# Structural basis of peptide recognition and activation of endothelin receptors

**DOI:** 10.1101/2023.01.17.524465

**Authors:** Yujie Ji, Jia Duan, Qingning Yuan, Xinheng He, Gong Yang, Shengnan Zhu, Kai Wu, Wen Hu, Tianyu Gao, Xi Cheng, Hualiang Jiang, H. Eric Xu, Yi Jiang

## Abstract

Endothelin system comprises three endogenous 21-amino-acid peptide ligands endothelin-1, -2, and -3 (ET-1/2/3), and two G protein-coupled receptor (GPCR) subtypes—endothelin receptor A (ET_A_R) and B (ET_B_R). Since ET-1, the first endothelin, was identified in 1988 as one of the most potent endothelial cell-derived vasoconstrictor peptides with long-lasting actions, the endothelin system has attracted extensive attention due to its critical role in vasoregulation and close relevance in cardiovascular-related diseases. Here we present three cryo-electron microscopy structures of ET_A_R and ET_B_R bound to ET-1 and ET_B_R bound to the selective peptide IRL1620. These structures reveal a highly conserved recognition mode of ET-1 and characterize the ligand selectivity by ETRs. They also present several conformation features of the active ETRs, thus revealing a specific activation mechanism. Together, these findings deepen our understanding of endothelin system regulation and offer an opportunity to design selective drugs targeting specific ETR subtypes.

## Introduction

Endothelin receptors (ETRs) is comprised by two subtypes (ET_A_R ^1^ and ET_B_R ^2^) with an approximately 60% sequence similarity, both of which belong to the class A G protein-coupled receptors (GPCRs) ^3^. ETRs are widely expressed in the human body, with ET_A_R primarily expressed in the cardiovascular system ^4^ and ET_B_R in the brain ^5^. They are activated by three types of endothelin (ET-1, ET-2, and ET-3), 21-amino-acid peptide hormones ^6,7^. Besides, naturally occurring sarafotoxins (s6a-d), isolated from snake venom, show a high degree of sequence similarity and biological effects with ET-1 (Fig. 1a) ^8^. Both ET_A_R and ET_B_R show different responses to these peptide agonists. ET_A_R exhibits similar subnanomolar affinities to ET-1 and ET-2 but a 100-fold weaker affinity to ET-3, while three ETs are equally effective for ET_B_R ^9,10^. Sarafotoxin s6c shows a 100-to 10,000-fold reduced affinity for ET_A_R compared with s6b, thus producing a substantial ET_B_R selectivity ^11^. Upon binding by agonists, ET_A_R primarily couples to the G_q_ protein, while ET_B_R couples to multiple G proteins, including G_s_, G_i_, and G_q_ proteins to transduce signals (Fig. 1b, data from GPCRdb.org ^12^). Intriguingly, two ETR subtypes exert opposing actions on vasoregulation: ET-1-mediated ET_A_R activation promotes long-lasting vasoconstriction, while ET_B_R mediates vasodilation and can inhibit the ET-1-mediated effects upon activated by ET-3, thus being considered physiological antagonist of ET_A_R ^13,14^.

**Fig. 1.**
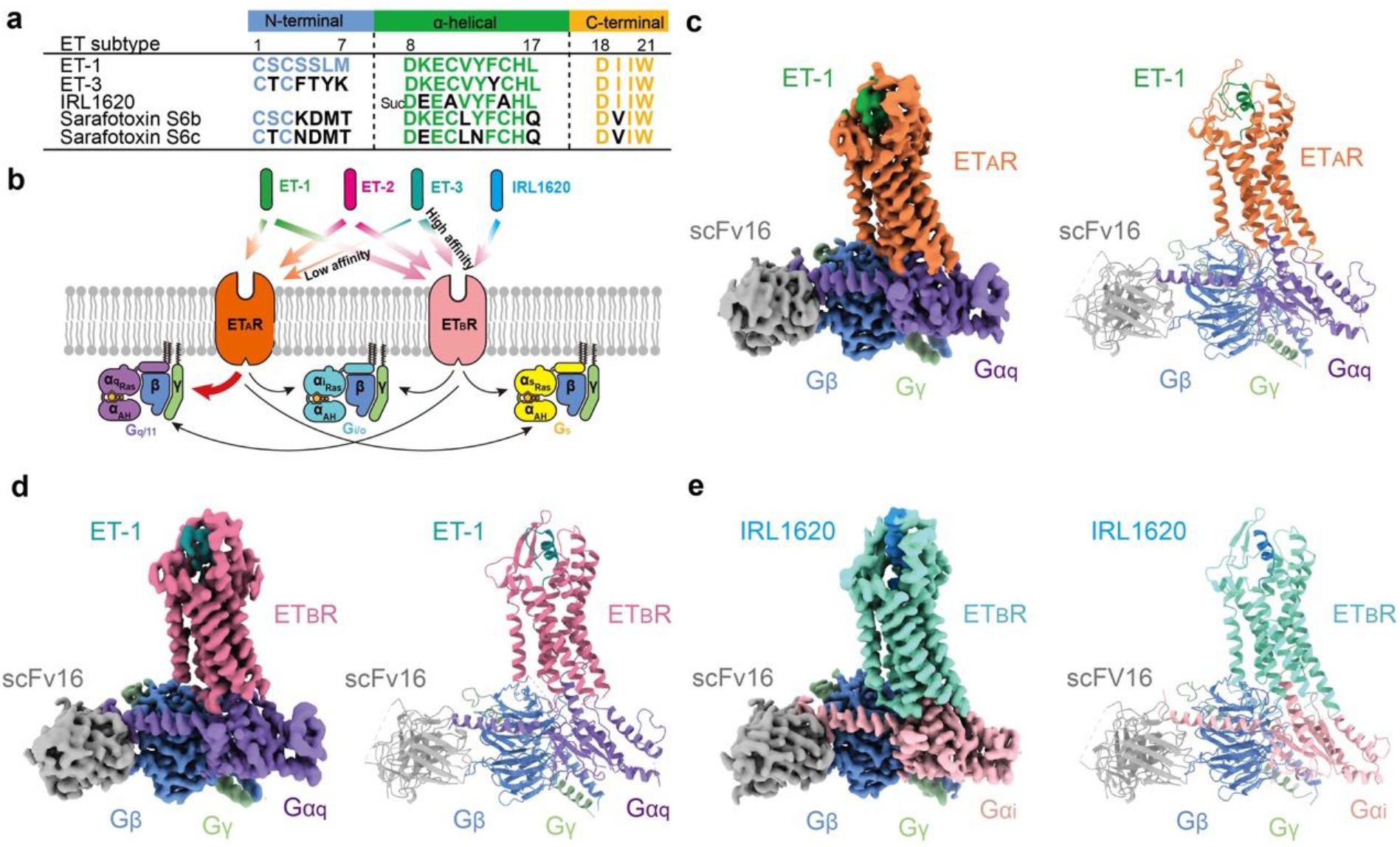
Cryo-EM structures of ET-1-ET_A_R-G_q_, ET-1-ET_B_R-G_q_ and IRL1620-ET_B_R-G_i_ complexes. **a** Sequence comparison of ET-1, ET-3, IRL1620, sarafotoxin S6b, and S6c. **b** Schematic illustration of G protein-coupling of ETRs activated by endothelin and IRL1620. **c-e** Cryo-EM density maps (left panel) and cartoon representation (right panel) of ET-1-ET_A_R-G_q_-scFv16 (**c**), ET-1-ET_B_R-G_q_-scFv16 (**d**), and IRL1620-ET_B_R-G_i_-scFv16 complexes (**e**). Components of ETRs complexes are colored as indicated.

The endothelin system has ignited extensive interest by its utmost importance in cardiovascular system regulation. Dysregulation of the endothelin system, especially the ET_A_R receptor, is closely associated with arterial hypertension, atherosclerosis, heart failure, renal disease, diabetes, etc., making ETRs promising drug targets for the treatment of these diseases ^9,10,14^. Several non-selective ETRs antagonists (bosentan and macitentan) and ET_A_R-selective antagonists (ambrisentan and clazosentan) haven approved for the treatment of pulmonary arterial hypertension (PAH). ET_B_R-selective agonist N-succinyl-[Glu^9^, Ala^11,15^]-ET-1(8-21), also named IRL1620 ^15^, is also in clinical trials for adjuvant cancer therapy to enhance the delivery of anti-tumor drugs by improving blood flow ^16^. Thus, developing selective ligands targeting specific endothelin receptor subtypes is of great pharmacology interest.

Extensive efforts have been made to clarify the recognition and activation mechanism of ETRs. Structural and relationship analysis had clarified residues on ET-1 crucial for binding ^17-19^. Crystal structures of ET_B_R bound to endogenous peptides, including ET-1, ET-3, s6b, and synthesized agonists or antagonists were subsequently released, thus offering molecular evidence on ET_B_R recognition by ligands and receptor activity regulation ^11,20-23^. In addition, difference architectures of ECL2 of ET_A_R and ET_B_R were suggested to account for selectivity of ET isopeptides ^22^. A structure-based molecular simulation revealed that E9 in sarafotoxin s6c may rearrange its interaction with extracellular regions, subsequently destabilize its interaction with ET_A_R, thus leading to the reduced ET_A_R affinity and ET_B_R specificity ^11^. However, the inaccessible structural information on ET_A_R subtype hampers our understanding of mechanisms of ligand recognition by ET_A_R, the ligand selectivity, and the ET_A_R activation. In this study, we present cryo-electron microscopy (cryo-EM) structures of engineered G protein-coupled ET_A_R and ET_B_R bound to ET-1 and ET_B_R bound to the selective peptide IRL1620. These structures reveal the molecular mechanism of peptide agonists recognition by ET_A_R and characterize the ligand-binding selectivity for both ETRs. These findings provide a structural template for the understanding of endothelin system regulation and offer an opportunity to design selective drugs targeting specific ETR subtypes.

## Results

### Structure determination of ET_A_R and ET_B_R complexes

The NanoBiT tethering strategy, which has been used to stabilize GPCR-G protein complex ^24^, was introduced to facilitate the assembly of three endothelin receptor complexes. The large subunit (LgBiT, 18 kDa) and a complimentary peptide (HiBiT, VSGWRLFKKIS) with high affinity for LgBiT were fused to C-termini of the receptor and the Gβ subunit, respectively (Supplementary information, Fig. S1). The complementation of LgBiT and HiBiT facilitates the stabilization of ETR and G protein. The chimeric mini-Gα_q_ used for the assembly of ETR complexes is the GTPase domain of the Gα_q_ subunit with its N-terminal 35 amino acids replaced by the corresponding sequence in Gα_i2_ to facilitate binding of a single-chain variable fragment (scFv16). It has been successfully applied in the structural determination of the 5-HT_2A_-Gα_q_ complex ^25^. The Gα_i_ subunit was engineered by introducing two dominant-negative mutations, G203A and A326S, to increase the stability of the ET_B_R-G_i_ protein complex ^26^. Unless otherwise specified, G_q_ and G_i_ refer to respective engineered G proteins, which are used in the structural studies of ETR complexes. The structure of ET-1 bound ET_A_R and ET_B_R in complex with engineered G_q_ heterotrimer were determined by single-particle cryo-EM at global resolutions of 3.0 Å and 3.5 Å. The cryo-EM structure of the engineered G_i_-coupled ET_B_R bound to the selective peptide IRL1620 was solved at a resolution of 3.0 Å (Fig. 1c-e; Supplementary information, Fig. S2 and Table S1). The high-quality map allowed accurate model building for receptor residues N66^NTD^ to N378^8.52^, I85^NTD^ to F397^8.54^, and T84^NTD^ to L401^8.58^ (superscripts refer to Ballesteros-Weinstein numbering ^27^) for the ET-1-ET_A_R-G_q_, ET-1-ET_B_R-G_q_, and IRL1620-ET_B_R-G_i_ complexes, respectively, which can provide detailed atomic information of the ligand-binding pocket and the receptor-G protein coupling interface (Fig. 1c-e; Supplementary information, Fig. S3a-c).

Globally, ET_A_R and ET_B_R adopt the canonical seven-transmembrane fold and G protein recruitment manner. All three complexes show a similar overall arrangement, with a root mean square deviation (RMSD) value for Cα atoms of 0.7-1.0 Å (Fig. 2a). At the receptor extracellular portion, both ETRs exhibit a large-scale outward kink at their extracellular end of TM5, which is distinguished by other class A peptide GPCRs. The kink is stabilized by a polar interaction network comprised of K^3.33^, E^4.60^, K^5.38^, Y^5.34^, and W21 of ET-1 and creates a wide-open pocket to accommodate peptide agonists (Fig. 2b, c). In addition, besides the conserved TM3-ECL2 disulfide bond, ETRs have an additional disulfide bond between C69/C90^NTD^ and C341/C358^7.25^ for ET_A_R/ET_B_R. The featured disulfide bond links receptor N-terminus and the extracellular end of TM7, thus capping the peptide-binding pocket. The compact extracellular interaction may prevent the agonist dissociation and explain the long-lasting activation of ET-1 (Fig. 2d).

**Fig. 2.**
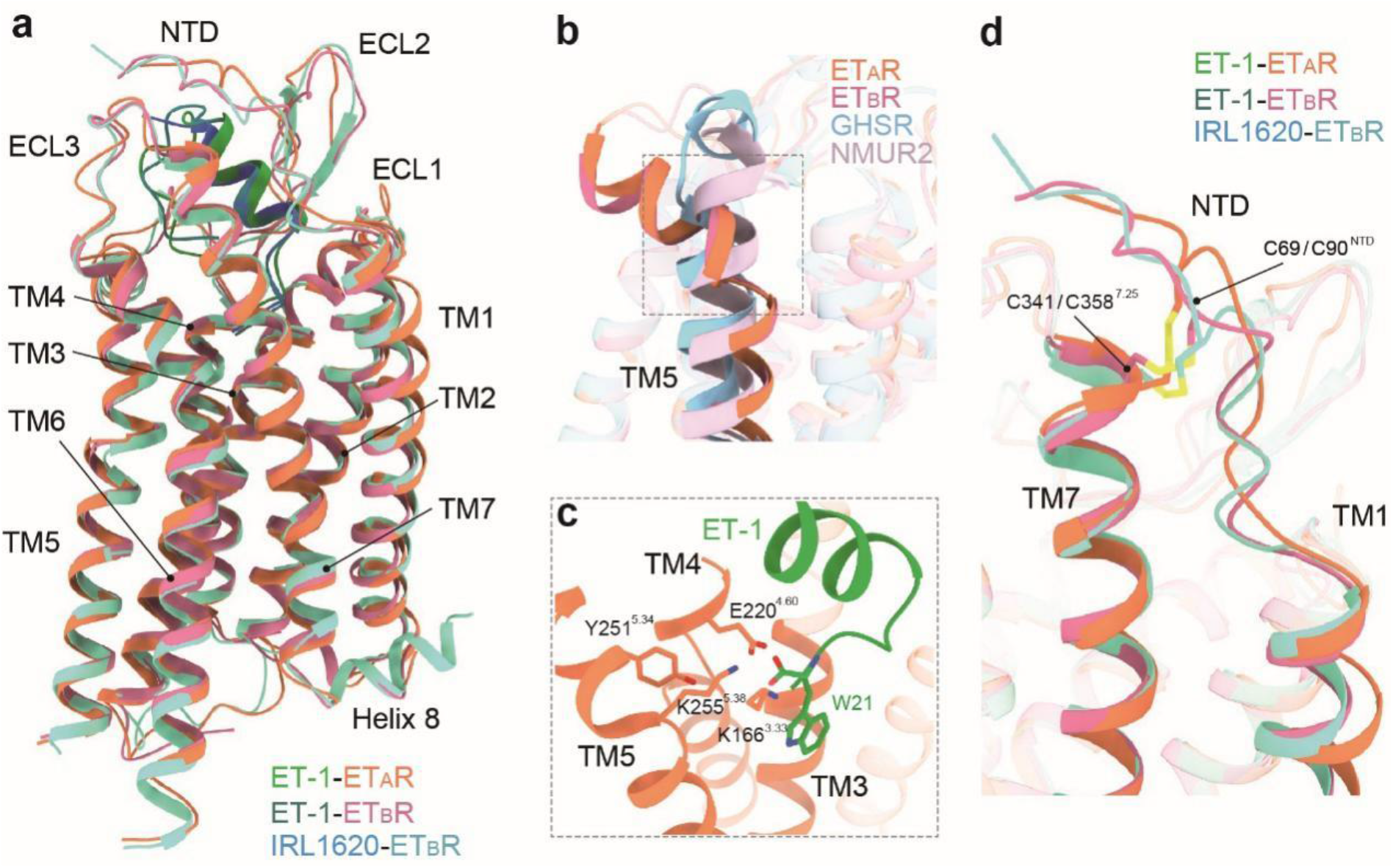
Structure features of endothelin receptors in the active state. **a** Structure superposition of ET1-bound and IRL1620-bound ETRs. **b** Structure superposition of ET-1-bound ETRs and other active class A peptide GPCRs, GHSR (PDB: 7F9Y) and NMUR2 (PDB: 7W55). The notable outward kink of the extracellular part of TM5 upon ETRs activation is highlighted. **c** Polar interaction network that stabilizes the kink conformation of TM5 of ET_A_R. **d** The disulfide bond between C69/C90^NTD^ and C341/C358^7.25^ for ET_A_R/ET_B_R.

### Recognition mechanism of ET-1 by ET_A_R

The ET-1 comprises the N-terminal (residues C1-M7), α-helical (D8-L17), and C-terminal regions (D18-W21) (Fig. 3a). It adopts a bicyclic architecture, formed by two pairs of intramolecular disulfide bonds (C1-C15 and C3-C11) (Fig. 3b), as observed in crystal and nuclear magnetic resonance (NMR) structures ^18,28^. A previous NMR study identified a stable conformation of the N-terminal and α-helical regions in solution, whereas the C-terminal region is flexible ^28^. The C-terminal region of ETs fixes its conformation by inserting into the receptor core. The extreme C-terminal amino acid W21 makes a H-bond with K166^3.33^ and faces hydrophobic residues V169^3.36^, L259^5.42^, W319^6.48^, L322^6.51^, and I355^7.39^. Its backbone CO also forms an H-bond with K166^3.33^. I19 and I20 are buried in a hydrophobic cavity comprising I136^2.60^, W146^ECL1^, P162^3.29^, F224^4.64^, Y352^7.36^, and I355^7.39^ (Fig. 3c; Supplementary information, Fig. S3d). In addition, D18 forms a salt bridge with R326^6.55^, which makes a polar contact with D351^7.35^ and further S325^6.54^, thus constituting an extensive polar interaction network with TM6 and TM7 of ET_A_R (Fig. 3d; Supplementary information, Fig. S3d). These structural observations are supported by the dramatically decreased ET-1 activity on alanine mutants of K166^3.33^, R326^6.55^, and D351^7.35^ (Supplementary information, Fig. S4a and Table S2). These findings are consistent with a previous report that the C-terminal region is essential for the potency of ET-1 ^29^, of which W21 shows critical roles in receptor binding and activation ^30^. Small molecular antagonists of ET_B_R bosentan and K8794 and reverse agonist IRL2500 mimic the hexapeptide of the C-terminal region ^21,23^, also indicating the importance of this segment in regulating receptor activity. Intriguingly, ET-3 and IRL1620 show an identical C-terminus with ET-1, while s6b and s6c bears a similar hydrophobic replacement from isoleucine to valine at position 19 relative to ET-1 (Fig. 1a). Correspondingly, C-terminus-interacting residues in both ETRs are conserved (Fig. 3e; Supplementary information, Fig. S5), proposing a conserved receptor-binding mode across these peptides.

**Fig. 3.**
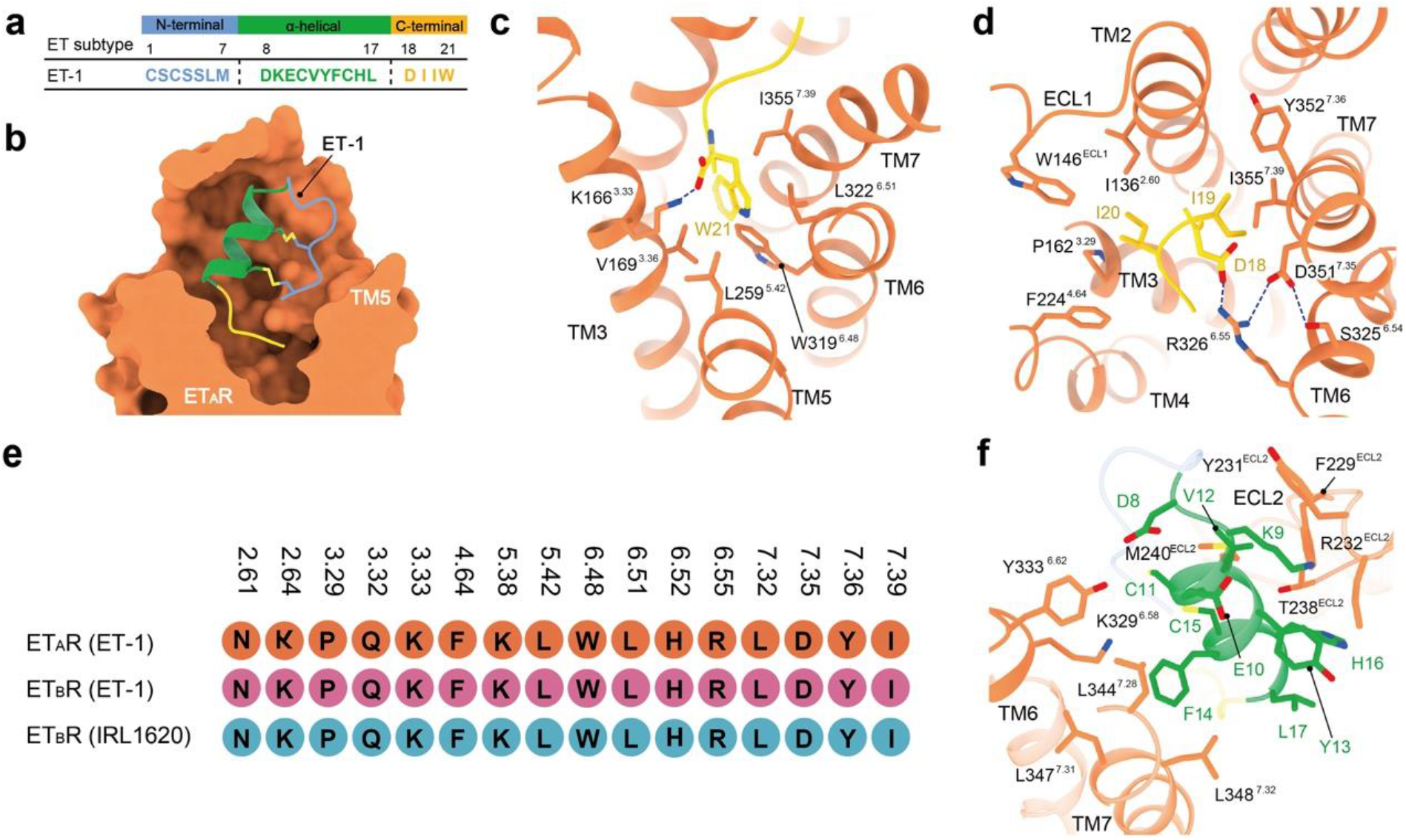
Recognition of ET-1 by ET_A_R. **a** Sequence of ET-1. The N-terminal, α-helical, and C-terminal regions of ET-1 are colored blue, green, and yellow, respectively. **b** Cross-section of the ET-1-binding pocket in ET_A_R. Two pairs of intramolecular disulfide bonds of ET-1 are displayed as sticks. ET-1 is shown as a ribbon model with its three regions colored as in **a. c-d** Detailed interactions between ET-1 and ET_A_R at the C-terminal region. H-bonds are shown as blue dotted lines. **e** The profile of the interaction residues between the C-terminal region of ligand and endothelin receptors. **f** Detailed interactions between ET-1 and ET_A_R at the α-helical region.

The N-terminal region of ET-1 is anchored to the α-helical region by two pairs of intramolecular disulfide bonds. Residues D8-L17 in ET-1 form a helical fold, constituting the architecture of the α-helical region (Fig. 3b). The side chain of D8 forms an H-bond with Y333^6.62^. The phenyl ring of F14 makes hydrophobic interactions with L344^7.28^, L347^7.31^, and L348^7.32^ (Fig. 3f). Residues on ECL2 of ET_A_R, including F229, Y231, R232, T238, and M240, are also critical for ET-1 activity through association with the α-helical region of the peptide (Supplementary information, Fig. S4a and Table S2). The N-terminal region of ET-1 points to the kinked N-terminus of receptor TM5 (Fig. 3b). Compared with the α-helical and C-terminal regions, the N-terminus of ET-1 exposes to the solvent with seldom ET_A_R residues making a minimal contribution to ET-1 activity. In contrast to the C-terminus region, the α-helical region, especially the N-terminus of peptide agonists, show lower sequence identity (Fig. 1a), indicating non-conserved binding modes of these two regions and their potential roles in regulating peptide selectivity against ET_A_R and ET_B_R.

### Ligand recognition selectivity for ET_A_R and ET_B_R

The structures of ET_A_R and ET_B_R provide templates to explore the mechanism of ligand-binding selectivity. Sequence comparison of ET-1 with two ET_B_R selective agonists, ET-3 and IRL1620, reveals that the most noticeable difference resides in the N-terminal region of peptides, in which all replaced amino acids in ET-3 from ET-1 are bulkier (Fig. 1a). This sequence difference elicits a hypothesis that ET_B_R has a larger pocket space to accommodate the bulkier N-terminal region of ET-3, which is supported by a smaller pocket size (1976 Å^3^, calculated by POVME ^31^) for ET_A_R relative to ET_B_R (2059 Å^3^). This conclusion is in agreement with a previous finding that increasing intramolecular ring size decreased peptide activity at ET_A_R and shifted the peptide towards ET_B_R selectivity ^32^.

Different from ETs, the N-terminal region is absent in agonist IRL1620 (Fig. 1a), which shows over 120,000-fold ET_B_R selectivity (*K*_i_ = 16 pM) over ET_A_R (*K*_i_ = 1900 nM) ^15^. Substituting the K9 of ET-1 with the glutamic acid (E9) in IRL1620 increases the electronegativity, which may facilitate the electrostatic interaction with the positively charged extracellular part of ET_B_R (Fig. 4a, b) ^22^. S6c bears an identical substitution from K9 to E9 compared with s6b (Fig. 1a). The charge of the 9^th^ residue has been thought to determine ETR selectivity of s6c over s6b ^33^, supporting the importance of electrostatic interaction on peptide selectivity. This theory is also consistent with a previous finding that the net negative electrostatic potential of D8-K9-E10 is essential for ETs binding to ET_B_R ^15^. In addition, the N^α^-succinylation of IRL1620 and its analogs increased the binding affinity for ET_B_R, but not ET_A_R, thus producing ET_B_R preference ^15^. The negative charge of succinylation is more compatible to the positively charged extracellular region of ET_B_R relative to its cognate receptor, probably contributing to the increased ET_B_R binding affinity and ET_B_R selectivity (Fig. 4b).

**Fig. 4.**
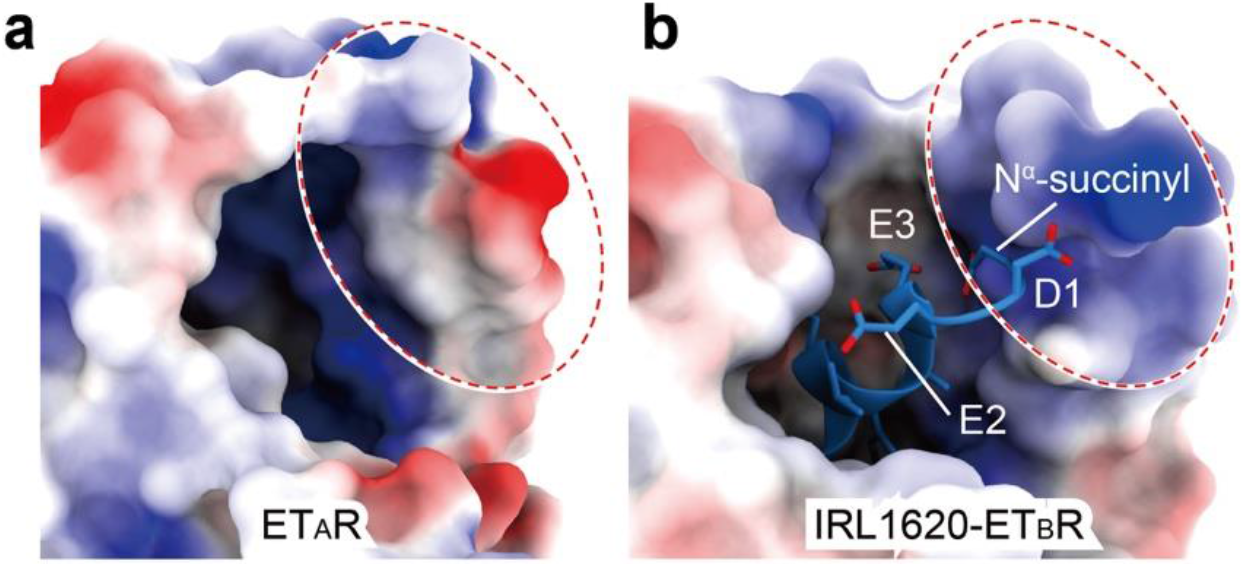
Electrostatic surfaces of the ET-1-bound ET_A_R and IRL1620-bound ET_B_R complex structures. The electrostatic surface presentation of the extracellular regions of ET-1 bound ET_A_R (**b**)and IRL1620 bound ET_B_R (**b**) are shown. ET-1 in the ET-1-ET_A_R complex structure is omitted for a clear presentation. The electrostatic potential was calculated and presented using PyMOL. Deep blue represents positively charged and red represents negatively charged areas.

### Activation of endothelin receptors

A structural comparison of ETR-G protein complexes with the crystal structures of ET_B_R in apo and agonist-, antagonist-, and reverse agonist-bound states reveals that these ETRs are indeed in the active state. TM6 of the ET-1 bound ET_B_R-G protein complex exhibits a notable outward displacement compared with that of crystal structures of apo (PDB: 5GLI, 4.4 Å, measured at Cα of R318^6.30^), antagonists/inverse agonist bound (PDB: 5XPR and 5×93, 5.4 and 4.6 Å), and the agonist ET-1 bound ET_B_R (PDB: 5GLH, 6.8 Å) due to the engagement of G proteins, the hallmark of class A GPCR activation (Fig. 5a). The almost identical active conformation of the ET_A_R relative to ET_B_R reveals a conserved activation mechanism in ETRs (Fig. 2a). Thus, structural comparison of the ET-1-ET_B_R-G_q_ complex with the apo ET_B_R was performed to analyze the activation mechanism of ETRs.

**Fig. 5.**
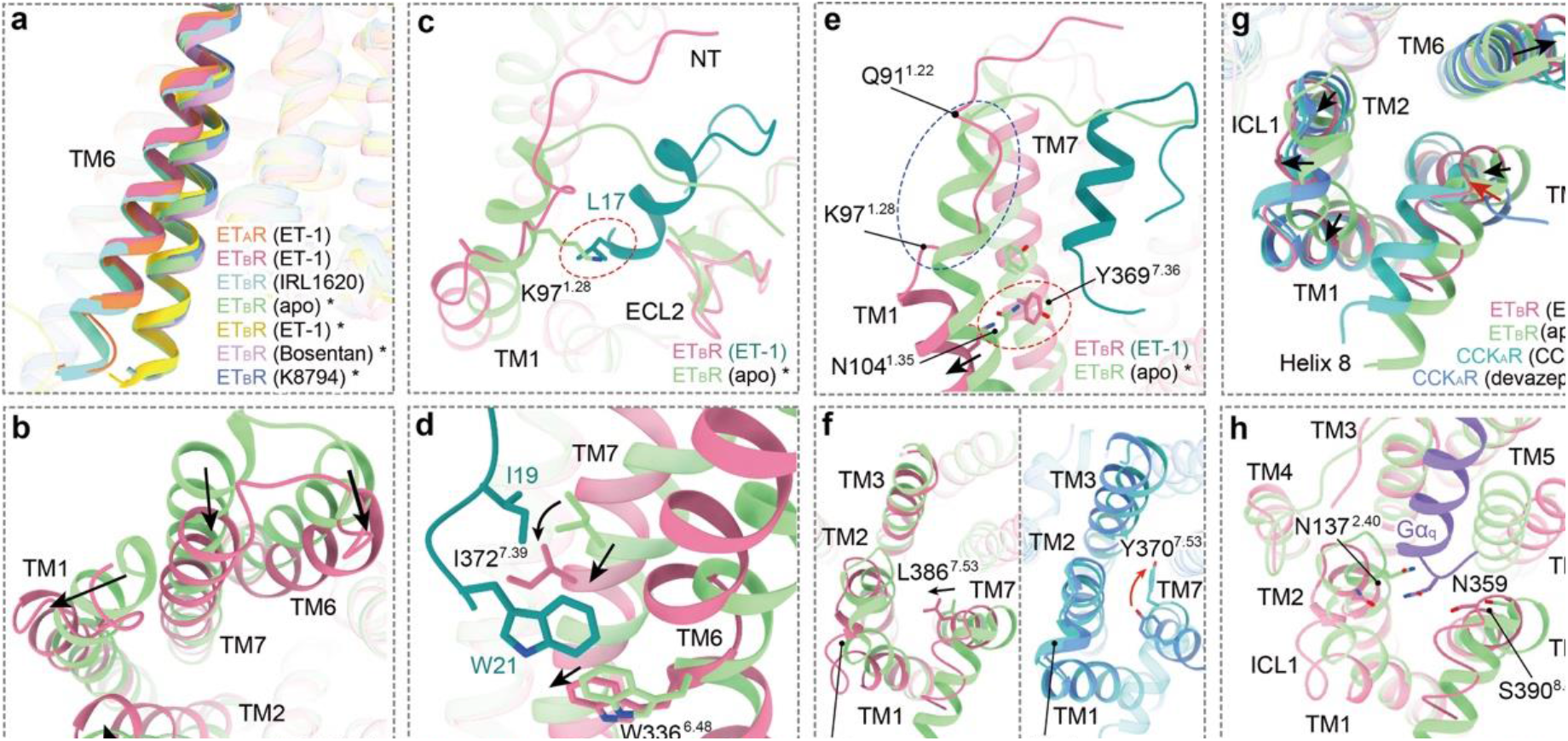
Activation and G protein-coupling of endothelin receptors. **a** Structure superposition of ETR-G protein complexes with the crystal structures of ET_B_R in apo (PDB: 5GLI) and agonist-(PDB: 5GLH), antagonist-(PDB: 5XPR), and reverse agonist-bound states (PDB: 5×93). **b** Overall conformational changes of the extracellular side of ET_B_R upon ET-1 binding. **c** The potential steric clash between L17 of ET-1 and K97^1.28^ of ET_B_R disturbs the receptor TM1 α-helix from Q91^1.22^ to K97^1.28^. The potential clash is highlighted by the red dashed oval. **d** Detailed Interaction between W21, I19 of ET-1 and W336^6.48^, I372^7.39^ of ET_B_R. The movement orientations of W366^6.48^, I372^7.39^, and TM7 upon ET_B_R activation are indicated as black arrows. **e** Conformation comparison of the extracellular part of TM1 between apo and ET-1 bound ET_B_R. The TM1 segment from Q91^1.22^ to K97^1.28^, highlighted in the blue dashed oval, undergoes a helix-to-loop conformational change upon ET_B_R activation. The clash between N104^1.35^ and Y369^7.36^ are indicated by a red dashed oval. **f** Differences between the conformational changes of N^7.49^P^7.50^xxY^7.53^ motif in ET_B_R and CCK_A_R. Y^7.53^ in the NPxxY motif is replaced by L^7.53^ in ETRs. The movement of L386^7.53^ in ET_B_R and Y370^7.53^ in CCK_A_R are indicated by black and red arrows, respectively. **g** Structure superposition of ET-1-bound ET_B_R, CCK-8-bound CCK_A_R (PDB: 7EZM), the crystal structures of ET_B_R in apo (PDB: 5GLI), and devazepide-bound CCK_A_R states (PDB: 7F8Y) in cytoplasmic view. Black arrows show the movement directions of the cytoplasmic ends of ET_B_R helices upon receptor activation, while the red arrow indicates the shift of TM7 upon CCK_A_R activation. **h** The polar interaction linking N359 of the Gα_q_ subunit and N137^2.40^ and S390^8.47^ of ET_B_R.

On the extracellular side, TM6 and TM7 of ET_B_R undergo inward movement upon ET-1 binding (3.7 and 4.8 Å, measured at Cα of Y350^6.62^ and R357^7.24^, respectively), compressing the ligand-binding pocket to form compact interactions with the peptide. Meanwhile, the extracellular end of TM2 occurs a laterally shift (2.9 Å, measured at Cα of A164^2.67^), while TM1 shows a smaller extent of inward bend (2.4 Å, measured at Cα of Q91^1.22^) and undergoes a helix-to-loop conformational change (Fig. 5b, c). The potential steric clash between L17 of ET1 and K97^1.28^ of ET^B^R disturbs the receptor TM1 α-helix from Q91^1.22^ to K97^1.28^ (Fig. 5c). The resulting loop approaches the center of the ligand-binding pocket and forms hydrophobic and van der Waal interactions with the α-helical region of ET-1. In addition, the α-helical region of ET-1 clashes with the N-terminus of ET_B_R in the apo state, disrupting the latter’s interaction with ECL2. Instead, the N-terminus of ET_B_R covers the ligand-binding pocket and form weak interactions with the N-terminus of the α-helical region of ET-1 (Fig. 5c). These movements of the receptor extracellular ends may propagate to the corresponding cytoplasmic end of the receptor to correspond with the seesaw model of class A GPCR activation, leading to the opening of the cytoplasmic cavity of the helical bundle to accommodate the G protein ^34^.

Insertion of ET-1 deep into the ligand-binding pocket leads to direct contact of W21 with the toggle switch residue W336^6.48^ in ET_B_R and triggers their downward movement. Intriguingly, the side chain of I372^7.39^ of ET_B_R undergoes a downward rotamer shift and a hydrophobic rearrangement with I19 of ET-1, leading to a half-helical downward movement of TM7 (Fig. 5d). In addition, the side chain of Y369^7.36^ occurs a remarkable rotation, which probably clashes with N104^1.35^ and pushes TM1 apart from the helical core (Fig. 5e).

The agonist binding propagates agonism signaling downward and triggers the rearrangement of conserved motifs, such as toggle switch, P^5.50^V^3.40^F^6.44^, and D^3.49^R^3.50^Y^3.51^ (Supplementary information, Fig. S6a-c). Intriguingly, Y^7.53^ in the conserved N^7.49^P^7.50^xxY^7.53^ motif is replaced by leucine in ETRs (Supplementary information, Fig. S6d). In the classic activation mechanism, the side chain of Y^7.53^ occurs a large amplitude of rotation towards TM3, causing the inward movement of TM7 and strengthening its packing with TM3 ^35^. Differently, upon ET_B_R activation, L386^7.53^ undergoes a lateral movement towards TM1 and TM2 (Fig. 5f), concomitant with the slight outward movement of these helices together with ICL1 (Fig. 5g), reflective of a unique activation mechanism of ETRs. This conformation may be further stabilized by the insertion of the α5 helix of the Gα subunit, in which N359 makes polar interactions with S390^8.47^ in the TM7-helix 8 hinge region and N137^2.40^ (Fig. 5h). Besides the abovementioned movements of helices, the downward propagated agonism signal eventually leads to the pronounced outward displacement of the cytoplasmic end of TM6, the hallmark of class A GPCRs activation, thus opening the cytoplasmic cavity of the receptor helix bundle to recruit G proteins (Fig. 5g).

## Discussion

Even though extensive efforts have been made, we are still far from fully understanding important issues in the regulation of the endothelin system, although several crystal structures of ET_B_R in the inactive state have been released. By determining cryo-EM structures of ET_A_R and ET_B_R in complex with peptide agonists and G proteins, here we provide molecular evidence to deepen our understanding of the rationale for ligand recognition, subtype selectivity, and activation of endothelin receptors. Our structures present several conformation features upon ETR activation, including the helix-to-loop change of the extracellular portion of TM5, upward half-helical, and shift towards TM1-ICL2-TM2 of TM7 due to unique inter-helical interactions (Fig. 2). Our structures also reveal a conserved recognition mechanism of endothelin isopeptides (ET-1/-2/-3) between both endothelin receptor subtypes (Fig. 3c-e). The identical C-terminus of ETs inserts deep into the ligand-binding pocket of ETRs and is critical for the ET-1-induced ETR activation. The extreme C-terminal residue W21 of ETs directly contacts with toggle switch residue W^6.48^ and triggers familial ETR activation (Fig. 3c). We further provide the structural explanations on peptide selectivity for ETRs, including differences between two ETRs in the pocket sizes and electrostatic potential of the receptor extracellular surface (Fig. 4), and also demonstrate the activation features of ETRs (Fig. 5). Thus, our findings offer a new opportunity for designing drugs targeting specific ETR subtypes for the treatment of diseases caused by endothelin system malfunction.

## Materials and Methods

### Cell culture

Spodoptera frugiperda 9 (*Sf9*) insect cells (Expression Systems) were grown in ESF 921 serum-free medium (Expression Systems) at 27°C and 120 rpm. AD293 cells were purchased from American Type Culture Collection (ATCC), cultured in Dulbecco’s modified Eagle’s medium (DMEM, Life Technologies) supplemented with 10% fetal bovine serum (FBS, Gibco), and maintained in a humidified chamber with 5% CO_2_ at 37°C.

### Constructs

The human wild-type (WT) ET_A_R and ET_B_R gene was codon-optimized for *Sf9* expression and synthesized by Synbio Technologies. To facilitate the expression and purification, human ET_A_R (residues 21-406) and human ET_B_R (residues 27-424) DNA was cloned into a modified pFastBac vector (Invitrogen), which contains an N-terminal FLAG tag (DYKDDDD) followed by an 8 × His-tag before the receptor using homologous recombination (CloneExpress One Step Cloning Kit, Vazyme). The native signal peptide of the receptors was replaced with the prolactin precursor sequence to increase protein expression.

NanoLuc Binary Technology (NanoBiT, Promega) is originally a structurally complementary reporter system to evaluate protein-protein interaction in cells ^36^. NanoBiT is composed of a large subunit (LgBiT, 18 kDa) and a 11 amino acid complimentary peptide (SmBiT, peptide, VTGYRLFEEIL) with low affinity for LgBiT, both of which are connected to two interacting target proteins of interest, respectively. When the two proteins interact, the two subunits assemble into an active enzyme and generate a bright luminescent signal in the presence of substrate. To improve the homogeneity and stability of ETR-G protein complexes, we applied the NanoBiT tethering strategy ^24^ by fusing a LgBiT subunit at the receptor C-terminus. Rat Gβ1 fuses a HiBiT subunit (peptide 86, VSGWRLFKKIS), a homolog peptide of SmBiT with high affinity for LgBiT ^36^, with a 15-amino acid (15AA) polypeptide linker (GSSGGGGSGGGGSSG) at its C-terminus (Supplementary information, Fig. S1).

The Gα_q_ was designed based on a miniGα_s_ skeleton with N-terminus replaced by Gα_i_ for the binding of scFv16. Bovine Gα_i_ were incorporated with two dominant-negative mutations (G203A and A326S) by site-directed mutagenesis to increase the stability of the Gαβγ complex and decrease the affinity of nucleotide-binding. The engineered Gα_q_, Gα_i_, Gβ1-15AA-HiBiT, and bovine Gγ2 were cloned into the pFastBac vector, respectively. scFv16 was cloned into a modified pFastBac vector which contains a GP67 secretion signal peptide at its N-terminus.

### Expression and purification of complexes

Baculoviruses were prepared using the Bac-to-Bac Baculovirus Expression System (Invitrogen). *Sf9* insect cells were cultured to a density of 3 × 10^6^ cells per mL and co-infected with FLAG-His 8-ET_A_R(21-406)/ET_B_R(27-424)-LgBiT, engineered Gα_q_/Gα_i_, Gβ1-15AA-HiBiT, Gγ2, and scFv16 baculoviruses at a 1:1:1:1:1 ratio. The cells were then harvested by centrifugation 48 h post-infection and stored in -80°C for future use.

The frozen cells were thawed on ice and resuspended in lysis buffer containing 20 mM HEPES, pH 7.5, 100 mM NaCl, 10% (v/v) glycerol, 10 mM MgCl_2_, 5 mM CaCl_2_, and supplemented with EDTA-free protease inhibitor cocktail (TargetMol). For ET-1-ET_A_R-G_q_/ IRL1620-ET_B_R-G_i_ complex, cells were lysed by dounce homogenization and complex formation was initiated with the addition of 25 mU/mL Apyrase (Sigma-Aldrich) and 10 μM ET-1/ IRL1620 (GenScript) for 1.5 h at room temperature (RT). The membrane was then solubilized by adding 0.5% (w/v) lauryl maltose neopentyl glycol (LMNG, Anatrace) and 0.1% (w/v) cholesterol hemisuccinate (CHS, Anatrace) for 2 h at 4°C. The sample was clarified by centrifugation at 30,000 rpm for 30 min and the supernatant was then incubated with anti-DYKDDDDK Affinity Beads (GenScript) for 3 h at 4°C. After incubation, the resin was collected by centrifugation (600 g, 10 min) and loaded into a gravity flow column, followed by wash with twenty-column volumes of 20 mM HEPES, pH 7.5, 100 mM NaCl, 10% (v/v) glycerol, 5 mM MgCl_2_, 1 μM ET-1/IRL1620, 0.01% (w/v) LMNG and 0.002% (w/v) CHS. The protein was finally eluted with ten-column volumes of 20 mM HEPES, pH 7.5, 100 mM NaCl, 10% (v/v) glycerol, 5 mM MgCl_2_, 1 μM ET-1/IRL1620, 0.01% (w/v) LMNG, and 0.2 mg/mL FLAG peptide. The purified complexes were concentrated using a Amicon Ultra centrifugal filter (molecular weight cut-off of 100 kDa, Millipore) and then subjected to a Superose 6 Increase 10/300 GL column (GE Healthcare) that was pre-equilibrated with buffer containing 20 mM HEPES, pH 7.5, 100 mM NaCl, 2 mM MgCl_2_, 1 μM ET-1/IRL1620, 0.00075% (w/v) LMNG, 0.00025% (w/v) GDN, and 0.0002% (w/v) CHS. The purification procedure of ET-1-ET_B_R-G_q_ complex was similar to ET_A_R, expect the addition of 0.05% (w/v) digitonin, the final size-column equilibrated with 0.05% (w/v) digitonin instead of 0.00075% (w/v) LMNG, 0.00025% (w/v) GDN and 0.0002% (w/v) CHS. The monomeric fractions of the complex were collected and concentrated for cryo-EM examination.

### Cryo-EM grid preparation and data collection

Three microliters of the purified ET-1-ET_A_R-G_q_-scFv16, ET-1-ET_B_R-G_q_-scFv16, and IRL1620-ET_B_R-G_i_-scFv16 complexes at the concentration of about 18 mg/mL, 10 mg/mL, and 20 mg/mL, respectively, were applied onto glow-discharged holey carbon grids (Quantifoil R1.2/1.3). Excess samples were blotted for 3s with a blot force of 1 and were vitrified by plunging into liquid ethane using a Vitrobot Mark IV (Thermo Fisher Scientific). Frozen grids were transferred to liquid nitrogen and stored for data acquisition. Cryo-EM imaging was performed on a Titan Krios G4 at 300 kV in the Advanced Center for Electron Microscopy at Shanghai Institute of Materia Medica, Chinese Academy of Sciences (Shanghai, China).

A total of 11,657 movies were collected for ET-1-ET_A_R-G_q_-scFv16 complex using a Gatan K3 detector in super-resolution mode with a pixel size of 0.824 Å using the EPU. Movies were obtained at a dose rate of about 15 electrons per Å^2^ per second with a defocus ranging from -0.8 to -1.8 μm. The total exposure time was 2.35 s, and 36 frames were recorded per micrograph.

For ET-1-ET_B_R-G_q_-scFv16 and IRL1620-ET_B_R-G_i_-scFv16 complexes, A total of 6,750 and 4,496 movies were collected by a Gatan K3 detector at a pixel size of 1.04 Å using the EPU, respectively. The micrographs were recorded in super-resolution mode at a dose rate of about 15 electrons per Å^2^ per second with a defocus ranging from -0.8 to -2 μm. The total exposure time was 3.13 s, and 36 frames were recorded per micrograph.

### Cryo-EM data processing

Cryo-EM image stacks were aligned using Relion 3.0 ^37^. Contrast transfer function (CTF) parameters for each micrograph were estimated by CTFFIND 4.0 ^38^. The following data processing was performed using RELION-4.0-beta ^37^.

For the ET-1-ET_A_R-G_q_-scFv16 complex, automated particle selection yielded 13,462,594 particle projections. The projections were subjected to reference-free 2D classification to discard poorly defined particles, producing 665,480 particle projections for further processing. This subset of particle projections was subjected to a round of maximum-likelihood-based three-dimensional classification with a pixel size of 0.824 Å. A selected subset containing 510,362 projections was used to obtain the final map using a pixel size of 0.824 Å. Further 3D classification without image alignment using a mask including only the receptor produced one good subset accounting for 510,197 particles, which were subsequently subjected to 3D refinement and Bayesian polishing. The final refinement generated a map with an indicated global resolution of 3.0 Å and was subsequently post-processed by DeepEMhancer.

For the ET-1-ET_B_R-G_q_-scFv16 complex, automated particle selection yielded 7,414,897 particle projections. The projections were subjected to reference-free 2D classification to discard poorly defined particles, producing 1,375,681 particle projections for further processing. This subset of particle projections was subjected to a round of maximum-likelihood-based three-dimensional classification with a pixel size of 1.04 Å. A selected subset containing 415,508 projections was used to obtain the final map using a pixel size of 1.04 Å. Further 3D classification focusing on the receptor without alignment using a mask to mask out the detergent and G protein produced one good subset accounting for 373,655 particles, which were subsequently subjected to 3D refinement, CTF refinement, and Bayesian polishing. The final refinement generated a map with an indicated global resolution of 3.5 Å, and was subsequently post-processed by DeepEMhancer.

For the IRL1620-ET_B_R-G_i_-scFv16 complex, automated particle selection yielded 4,078,200 particle projections. The projections were subjected to reference-free 2D classification to discard poorly defined particles, producing 967,642 particle projections for further processing. This subset of particle projections was subjected to a round of maximum-likelihood-based three-dimensional classification with a pixel size of 1.04 Å. A selected subset containing 410,751 projections was used to obtain the final map using a pixel size of 1.04 Å. Further 3D classification focusing the alignment on the receptor produced one good subset accounting for 191,493 particles, which were subsequently subjected to 3D refinement, CTF refinement, and Bayesian polishing. The final refinement generated a map with an indicated global resolution of 3.0 Å.

### Model building and refinement

The initial templates of ET_A_R and ET_B_R were derived from Alphafold2 ^39^. Models were docked into the EM density map using UCSF Chimera ^40^. The initial models were then subjected to iterative rounds of manual adjustment based on the side-chain densities of bulky aromatic amino acids in Coot ^41^ and automated refinement in Rosetta and PHENIX ^42^. The final refinement statistics were validated using the module “comprehensive validation (cryo-EM)” in PHENIX ^42^. The final refinement statistics are provided in Supplementary information, Table S1. All structural figures were prepared using Chimera^40^, Chimera X^43^, and PyMOL (https://pymol.org/2/).

### NanoBiT assay for β-arrestin recruitment

The NanoBiT assay was used to assess the interaction between ETRs and β-arrestin, which reflects the activity of β-arrestin recruitment by ETRs. Briefly, the full-length ET_A_R/ET_B_R and mutants were cloned into pBiT1.1 vector (Invitrogen) with a FLAG tag at its N-terminus and the LgBiT at its C-terminus. Human β-arrestin 2 (residues 1-393, with I386A, V387A, and F388A mutations) was cloned into pBiT2.1 vector (Invitrogen) with the SmBiT at its N-terminus. AD293 cells were cultured in DMEM/high Glucose (GE healthcare) supplemented with 10% (w/v) FBS (Gemini). Cells were maintained at 37 °C in a 5% CO_2_ incubator with 300,000 cells per well in a 6-well plate. Cells were grown overnight and then transfected with 1.5 μg ET_A_R/ET_B_R and 1.5 μg β-arrestin constructs by FuGENE® HD transfection reagent in each well for 24 h. Cells were harvested and re-suspended in Hanks’ balanced salt solution buffer (HBSS) at a density of 5 × 10^5^ cells/ml. The cell suspension was seeded in a 384-well plate at a volume of 10 μl per well, followed by 10 μl HBSS, 10 μl HBSS containing different concentrations of ET-1, and another 10 μl the NanoLuc substrate (furimazine, 1:25 dilution, Promega) diluted in the detection buffer. The luminescence signal was measured with an EnVision plate reader at room temperature.

### Receptor surface expression

Cell-surface expression levels of WT or mutants ET_A_R/ET_B_R were quantified by flow cytometry. AD 293 cells were seeded at a density of 1.5 × 10^5^ per well into 12-well culture plates. Cells were grown overnight and then transfected with 1.0 μg ET_A_R/ET_B_R construct by FuGENE® HD transfection reagent in each well for 24 h. After 24 h of transfection, cells were washed once with PBS and then detached with 0.2% (w/v) EDTA in PBS. Cells were blocked with PBS containing 5% (w/v) BSA for 15 min at room temperature before incubating with primary anti-Flag antibody (diluted with PBS containing 5% BSA at a ratio of 1:300, Sigma) for 1 h at room temperature. Cells were then washed three times with PBS containing 1% (w/v) BSA and then incubated with anti-mouse Alexa-488-conjugated secondary antibody (diluted at a ratio of 1:1000, Invitrogen) at 4 °C in the dark for 1 h. After another three times of washing, cells were collected, and fluorescence intensity was quantified in a BD Accuri C6 flow cytometer system (BD Biosciences) through a BD Accuri C6 software1.0.264.21 at excitation 488 nm and emission 519 nm. Approximately 10,000 cellular events per sample were collected, and data were normalized to the wild-type ET_A_R/ET_B_R. Experiments were performed at least three times, and data were presented as means ± S.E.M.

### Statistical analysis

All functional study data were analyzed using Prism 8 (GraphPad) and presented as means ± S.E.M. from at least three independent experiments. Concentration-response curves were evaluated with a three-parameter logistic equation. *EC*_*50*_ is calculated with the Sigmoid three-parameter equation.

### Data availability

Density maps and structure coordinates have been deposited in the Electron Microscopy Data Bank (EMDB) and the Protein Data Bank (PDB) with accession codes EMD-34663 and 8HCQ for the ET-1-ET_A_R-G_q_-scFv16 complex; EMD-34667 and 8HCX for the ET-1-ET_B_R-G_q_-scFv16 complex; EMD-34619 and 8HBD for the IRL1620-ET_B_R-G_i_-scFv16 complex.

## Acknowledgments

The Cryo-EM data of the ET-1-ET_A_R-G_q_-scFv16, ET-1-ET_B_R-G_q_-scFv16, and IRL1620-ET_B_R-G_i_-scFv16 complexes were collected at the Advanced Center for Electron Microscopy, Shanghai Institute of Materia Medica (SIMM). We thank all the staff at the cryo-EM facilities for their technical support. This work was partially supported by the National Natural Science Foundation (32171187 to Y.J., 82121005 to Y.J., and H.E.X., 32130022 to H.E.X.); the Ministry of Science and Technology (China) grants (2018YFA0507002 to H.E.X.); the Shanghai Municipal Science and Technology Major Project (2019SHZDZX02 to H.E.X.); Shanghai Municipal Science and Technology Major Project (H.E.X.); the CAS Strategic Priority Research Program (XDB37030103 to H.E.X.).

## Author contribution

Y-J.J designed the expression constructs, purified the ETR proteins, prepared the final samples for negative stain and cryo-EM data collection, conducted functional studies, and participated in figure and manuscript preparation with the help of J.D.; Q.Y. performed cryo-EM data calculations, model building, and participated in supplementary figure preparation; S.Z. and T.G. participated in protein expression and functional studies; X.H. participated in figure preparation; H.J. and X.C. supervised

X.H. in figure preparation; K.W. and W.H. participated in cryo-EM grid preparation and data collection; Y.J. and H.E.X. conceived and supervised the project, analyzed the structures, and wrote the manuscript with inputs from all authors.

## Supplementary information, Figures

**Fig. S1.**
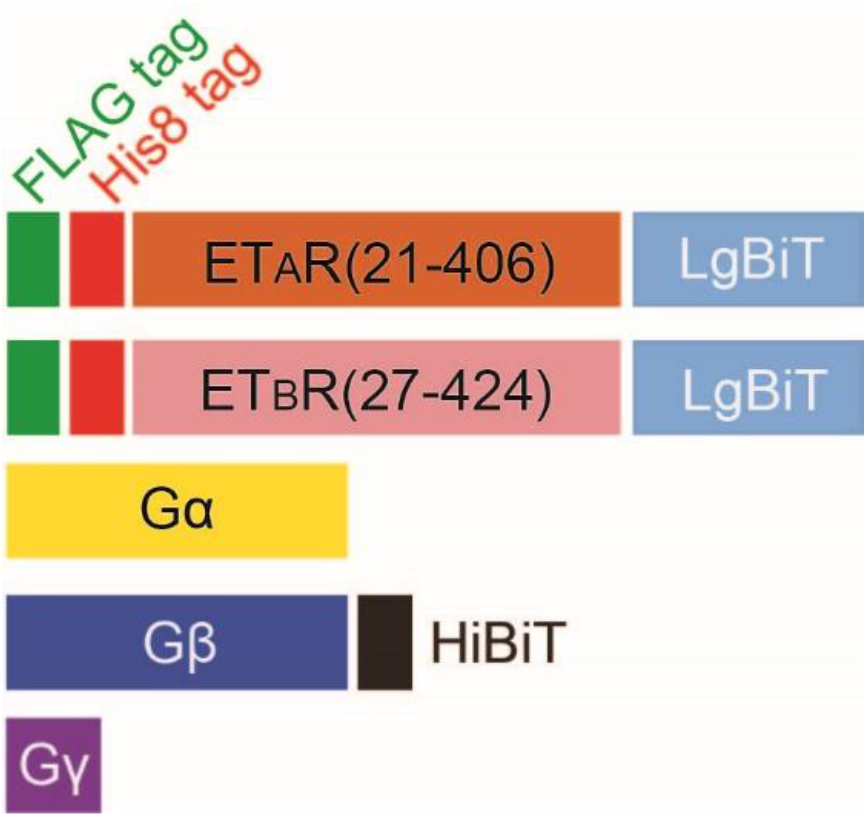
Engineered constructs used for expression of ET_A_R/ET_B_R-G protein complexes.

**Fig. S2.**
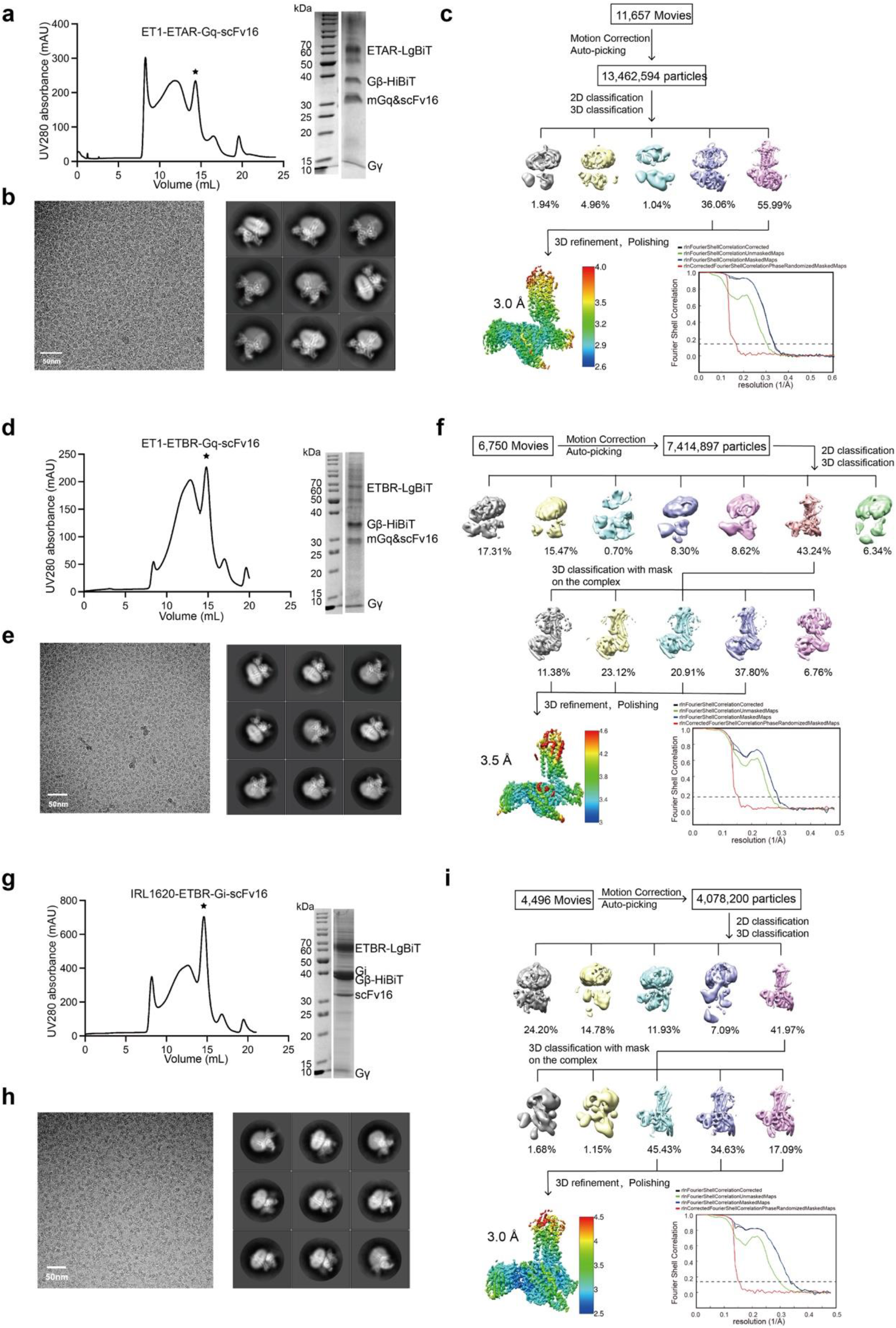
The ET-1-ET_A_R-G_q_-scFv16, ET-1-ET_B_R-G_q_-scFv16, and IRL1620-ET_B_R-G_i_-scFv16 complexes purification and cryo-EM data processing. **a, d, g** Representative elution profile and SDS-PAGE analysis of the ET-1-ET_A_R-G_q_-scFv16 (**a**), ET-1-ET_B_R-G_q_-scFv16 (**d**) and IRL1620-ET_B_R-G_i_-scFv16 complexes (**g**). Black asterisks denote the monomer of three complexes. **b, e, h** Cryo-EM micrographs and reference-free 2D class averages of the ET-1-ET_A_R-G_q_-scFv16 (**b**), ET-1-ET_B_R-G_q_-scFv16 (**e**) and IRL1620-ET_B_R-G_i_-scFv16 complexes (**h**). **c, f, i** Flow chart of the cryo-EM data processing for the ET-1-ET_A_R-G_q_-scFv16 (**c**), ET-1-ET_B_R-G_q_-scFv16 (**f**) and IRL1620-ET_B_R-G_i_-scFv16 complexes (**i**). The “Gold-standard” Fourier shell correlation (FSC) curve indicates that the overall resolution of the electron density map of the ET-1-ET_A_R-G_q_-scFv16 complex is 3.0 Å, the ET-1-ET_B_R-G_q_-scFv16 complex is 3.5 Å, and the IRL1620-ET_B_R-G_i_-scFv16 complex is 3.0 Å.

**Fig. S3.**
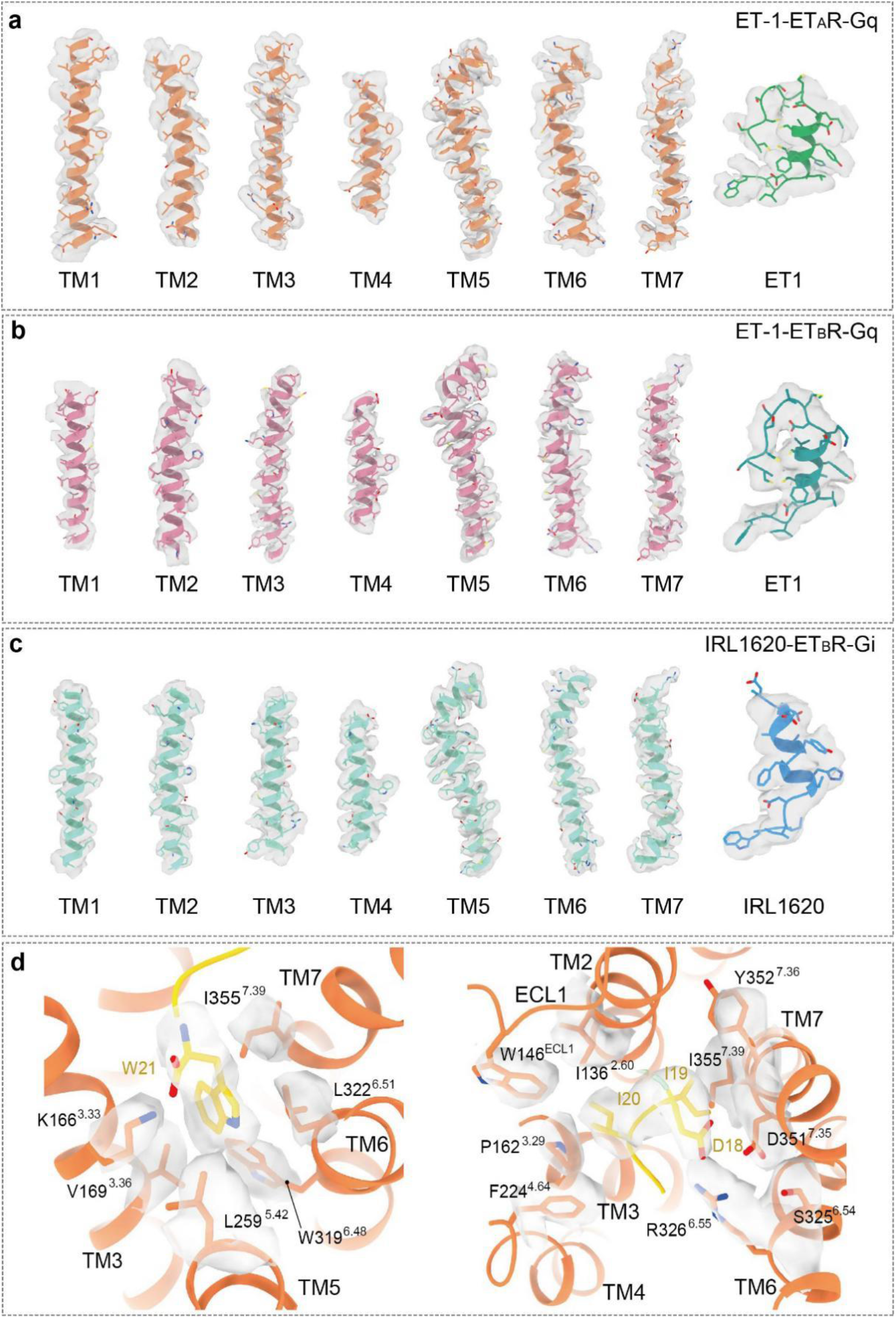
Representative EM density and coordinate of the ET-1-ET_A_R-G_q_-scFv16, ET-1-ET_B_R-G_q_-scFv16 and IRL1620-ET_B_R-G_i_-scFv16 complexes. EM density and model of ET-1-ET_A_R-G_q_ (**a**), ET-1-ET_B_R-G_q_ (**b**), IRL1620-ET_B_R-G_i_ complexes (**c**), and key ET_A_R residues in the binding site of the ET-1 C-terminus (**d**).

**Fig. S4.**
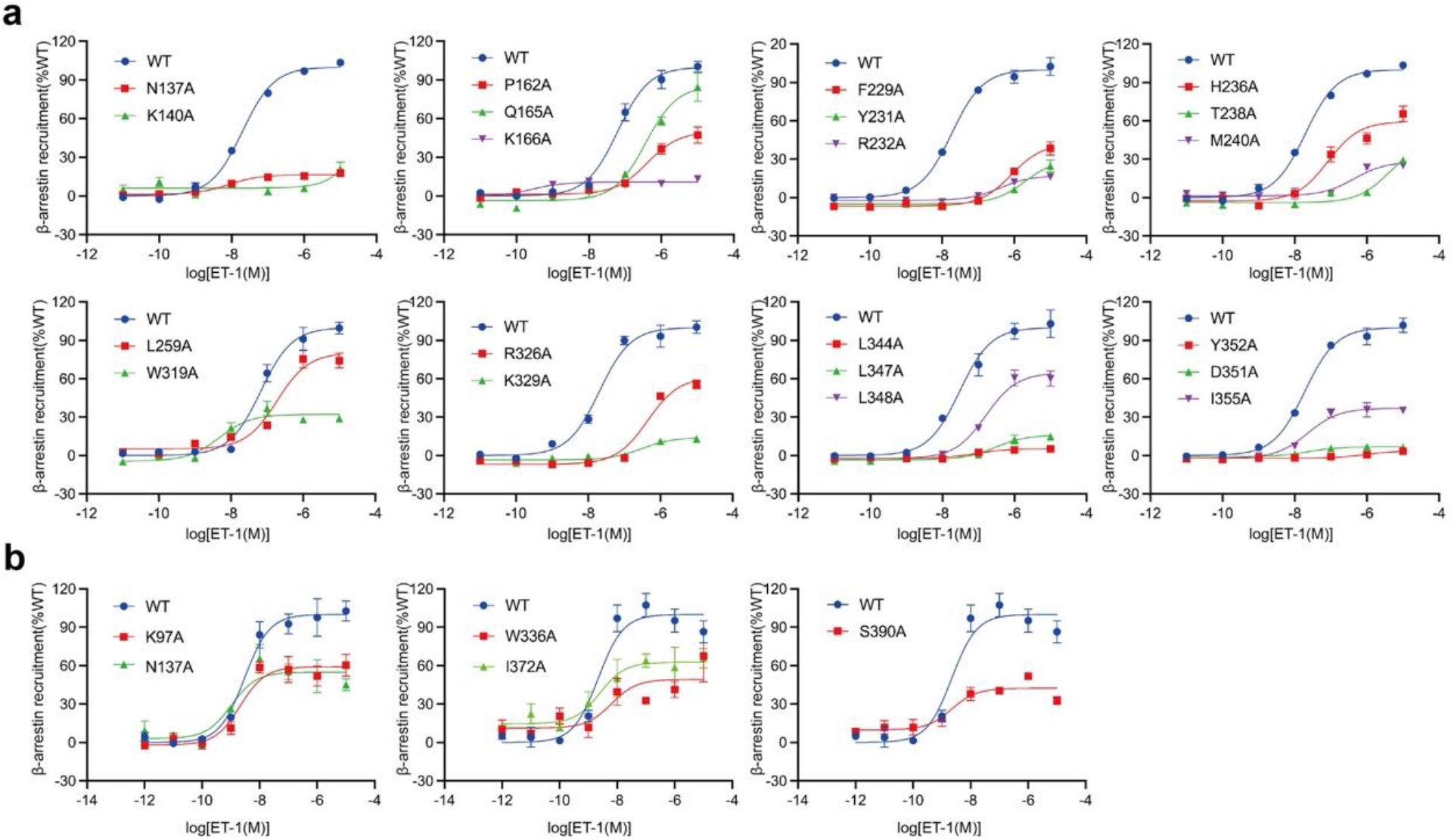
ET-1 response curves on WT and mutant ET_A_R/ET_B_R. WT or mutant ET_A_R (**a**) or ET_B_R (**b**) were transfected into AD293 cells and β-arrestin recruitment signals were measured to reflect the activity of ET-1. The response data was normalized by WT receptor within each individual experiment, with the basal activity for WT as 0, while the fitted *E*_*max*_ of WT as 100. Each point represents mean ± S.E.M. from three independent experiments. The representative concentration-response curves were shown.

**Fig. S5.**
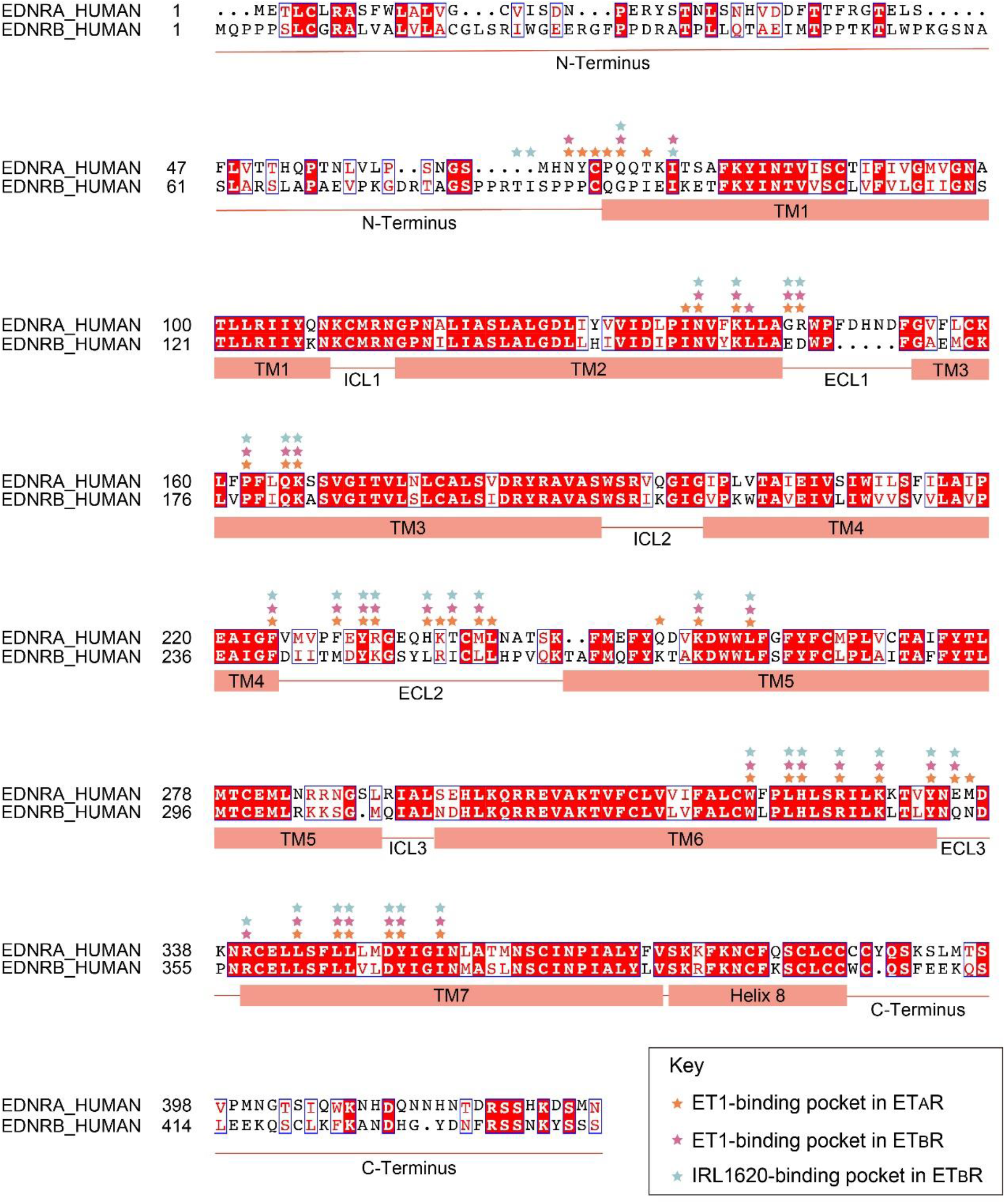
Sequence alignment of the ETRs. The sequence alignment of ET_A_R and ET_B_R was generated using NCBI and the graphics was created on the ESPript 3.0 server. α-helices for ETRs are shown as columns underneath the sequences. Orange asterisks represent the binding-pocket residues of ET_A_R bound to ET-1, pink asterisks represent the binding-pocket residues of ET_B_R bound to ET-1 and cyan asterisks represent the binding-pocket residues of ET_B_R bound to IRL1620.

**Fig. S6.**
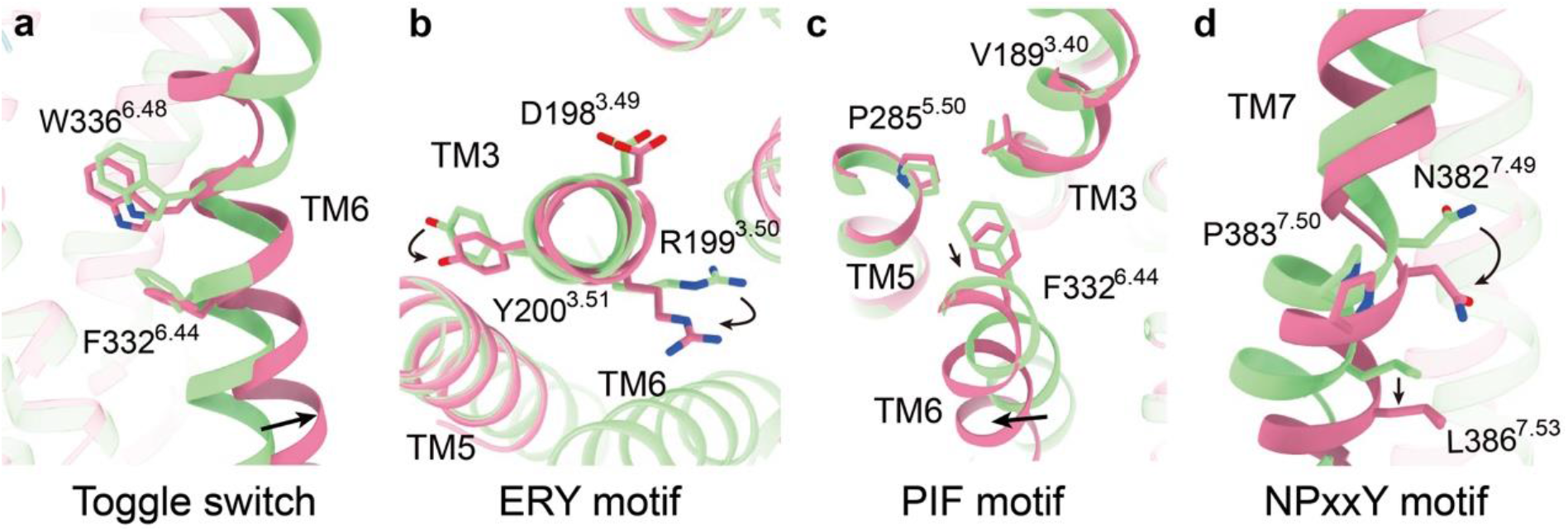
Conformational changes of the conserved “micro-switches” upon receptor activation. Toggle switch (**a**), ERY motif (**b**), PIF motif (**c**), NPxxY motif (**d**). The outward movement of TM6 of the active receptor is highlighted as a black arrow (**a**). The conformational changes of residue side chains are highlighted as black arrows.

**Table S1.** Cryo-EM data collection, model refinement, and validation statistics.

|  | ET-1-ET<sub>A</sub>R-G<sub>q</sub>-<br>scFv16 | ET-1-ET<sub>B</sub>R-G<sub>q</sub>-<br>scFv16 | IRL1620-ET<sub>B</sub>R-G<sub>i</sub>-<br>scFv16 |
| --- | --- | --- | --- |
| **Data collection and processing** |  |  |  |
| Magnification | 105K | 81K | 81K |
| Voltage (kV) | 300 | 300 | 300 |
| Electron exposure (e<sup>-</sup>/Å<sup>2</sup>) | 50 | 50 | 50 |
| Defocus range (μm) | -0.8 ~ -1.8 | -0.8 ~ -2 | -0.8 ~ -2 |
| Pixel size (Å) | 0.824 | 1.04 | 1.04 |
| Symmetry imposed | C1 | C1 | C1 |
| Initial particle projections (no.) | 13,462,594 | 7,414,897 | 4,078,200 |
| Final particle projections (no.) | 510,197 | 373,655 | 191,493 |
| Map resolution (Å) | 3.0 | 3.5 | 3.0 |
| FSC threshold | 0.143 | 0.143 | 0.143 |
| Map resolution range (Å) | 2.6-4.0 | 3.0-4.6 | 2.5-4.5 |
| **Refinement** |  |  |  |
| Refinement package | PHENIX-1.17.1-3660 | PHENIX-1.17.1-3660 | PHENIX-1.17.1-3660 |
| Real or reciprocal space | Real space | Real space | Real space |
| Model-Map CC (mask) | 0.78 | 0.7 | 0.76 |
| Model resolution (Å) | 3.4 | 4.0 | 3.4 |
| FSC threshold | 0.5 | 0.5 | 0.5 |
| B factors (Å<sup>2</sup>, mean value) |  |  |  |
| Protein residues | 93.41 | 82.66 | 84.79 |
| Ligands |  |  |  |
| **Model composition** |  |  |  |
| Non-hydrogen atoms | 9064 | 8946 | 8867 |
| Protein residues | 1157 | 1153 | 1155 |
| R.m.s. deviations |  |  |  |
| Bond lengths (Å) | 0.002 (0) | 0.002 (0) | 0.003 |
| Bond angles (°) | 0.609 (4) | 0.622 (1) | 0.603 |
| **Validation** |  |  |  |
| MolProbity score | 1.67 | 1.61 | 1.59 |
| Clashscore | 9.44 | 11.66 | 9.16 |
| Rotamer outliers (%) | 0.41 | 0.52 | 0.53 |
| **Ramachandran plot** |  |  |  |
| Favored (%) | 96.99 | 97.88 | 97.52 |
| Allowed (%) | 2.83 | 2.03 | 2.39 |
| Disallowed (%) | 0.18 | 0.09 | 0.09 |
| **Data availability** |  |  |  |
| PDB entry | 8HCQ | 8HCX | 8HBD |
| EMDB entry | EMD-34663 | EMD-34667 | EMD-34619 |

**Table S2.** Effects of ET-1 on ET_A_R/ET_B_R mutants. NanoBiT assay was performed to evaluate the effects of ET1 on β-arrestin 2 of ET_A_R/ET_B_R. The surface expression of each ET_A_R/ET_B_R mutant was normalized to wild-type (WT) receptor, which was set to 100%. Data are presented as means ± S.E.M. of three independent experiments in triplicate (n = 3). N.D., not determined.

| Receptor | Residue number | Mutant | *EC*<sub>50</sub> (nM) | Fold of change | Surface expression(%WT) |
| --- | --- | --- | --- | --- | --- |
| ET<sub>A</sub>R | - | WT | 33.73±8.05 | 1 | 100 |
| | 2.61 | N137A | N.D. | - | 87.85±2.65 |
| | 2.64 | K140A | N.D. | - | 88.94±6.76 |
| | 3.29 | P162A | >376.50 | >11.16 | 97.65±6.94 |
| | 3.32 | Q165A | >305.30 | >9.05 | 63±1.3 |
| | 3.33 | K166A | N.D. | - | 126.09±7.12 |
| | ECL2 | F229A | >657.60 | >19.50 | 105.28±7.64 |
| | ECL2 | Y231A | N.D. | - | 64.39±7.04 |
| | ECL2 | R232A | N.D. | - | 80.72±8.26 |
| | ECL2 | H236A | 60.92±8.39 | 1.81 | 91.01±8.39 |
| | ECL2 | T238A | N.D. | - | 55.85±5.13 |
| | ECL2 | M240A | N.D. | - | 72.84±4.46 |
| | 5.42 | L259A | 190.73±11.26 | 5.66 | 60.26±9.19 |
| | 6.48 | W319A | 5.89±0.83 | 0.17 | 108.92±2.68 |
| | 6.55 | R326A | >463.50 | >13.74 | 60.86±8.38 |
| | 6.58 | K329A | N.D. | - | 88.41±5.28 |
| | 7.28 | L344A | 127.96±17.23 | 3.79 | 100.89±8.23 |
| | 7.31 | L347A | N.D. | - | 69.16±3.76 |
| | 7.32 | L348A | N.D. | - | 70.72±0.43 |
| | 7.35 | D351A | N.D. | - | 106.9±2.04 |
| | 7.36 | Y352A | N.D. | - | 80.24±6.83 |
| | 7.39 | I355A | N.D. | - | 54.53±6.4 |
| ET<sub>B</sub>R | - | WT | 2.46±0.18 | 1 | 100 |
| | 1.28 | K97A | 1.68±0.33 | 0.69 | 85.63±2.04 |
| | 2.40 | N137A | 2.79±0.88 | 1.14 | 50.48±4.73 |
| | 6.48 | W336A | 5.55±0.58 | 2.26 | 71.82±4.97 |
| | 7.39 | I372A | 2.12±0.34 | 0.86 | 111.70±5.32 |
| | 8.47 | S390A | 2.57±0.48 | 1.05 | 110.75±3.33 |

